# Remodeling of pre-existing myelinated axons and oligodendrocyte differentiation is stimulated by environmental enrichment in the young adult brain

**DOI:** 10.1101/2020.01.21.914572

**Authors:** Madeline Nicholson, Rhiannon J Wood, Jessica L Fletcher, David G Gonsalvez, Anthony J Hannan, Simon S Murray, Junhua Xiao

## Abstract

Oligodendrocyte production and central nervous system (CNS) myelination is a protracted process, extending into adulthood. While stimulation of neuronal circuits has been shown to enhance oligodendrocyte production and myelination during development, the extent to which physiological stimuli induces activity-dependent plasticity within oligodendrocytes and myelin is unclear, particularly in the adult CNS. Here, we find that using environmental enrichment (EE) to physiologically stimulate neuronal activity for 6-weeks during young adulthood in C57Bl/6 mice results in an enlargement of callosal axon diameters, with a corresponding increase in thickness of pre-existing myelin sheaths. Additionally, EE uniformly promotes the direct differentiation of pre-existing oligodendroglia in both corpus callosum and somatosensory cortex, while differentially impeding OPC homeostasis in these regions. Furthermore, results of this study indicate that physiologically relevant stimulation in young adulthood exerts little influence upon the *de novo* generation of new myelin sheaths on previously unmyelinated segments and does not enhance OPC proliferation. Rather in this context, activity-dependent plasticity involves the coincident structural remodeling of axons and pre-existing myelin sheaths and increases the direct differentiation of pre-existing oligodendroglia, implying constraints on maximal *de novo* production in the adult CNS. Together, our findings of myelinated axon remodeling and increased pre-existing oligodendroglial differentiation constitute a previously undescribed form of adaptive myelination that likely contributes to neuronal circuit maturation and the maintenance of optimum cognitive function in the young adult CNS.

**Main points:**

- Environmental enrichment induces the plasticity of myelinated axons, resulting in axon caliber enlargement and increased thickness of pre-existing myelin sheaths
- Environmental enrichment increases the direct differentiation of pre-existing oligodendroglia
- Environmental enrichment alters OPC homeostasis

## Introduction

Oligodendrocytes (OLs) are critical components of the central nervous system (CNS), both generating the insulating myelin sheaths that facilitate rapid transmission of action potentials along axons and providing essential metabolic and trophic support to neurons (Nave & Werner, 2014). Myelination begins developmentally and is a lifelong process, with the continued production of new OLs and generation of new myelin sheaths occurring up until middle-age (Hill, Li, & Grutzendler, 2018). Studies have revealed that the myelinating process is receptive to neuronal activity (Foster, Bujalka, & Emery, 2019) and activity-dependent plasticity during early postnatal development increases oligodendrogenesis and myelin production, with a myelination bias towards selectively activated axons ((Gibson et al., 2014; Stanislaw Mitew et al., 2018). Conversely, downregulations in activity via social isolation during early postnatal development decreases the number of internodes and thickness of myelin sheaths (Liu et al., 2012; Makinodan, Rosen, Ito, & Corfas, 2012). These experimental findings, along with computational evidence that myelination pattern variations (e.g. changes to the number, length or thickness of sheaths) can markedly alter nerve conduction velocity (Castelfranco & Hartline, 2015), identifies a role for myelin in fine-tuning neuronal connectivity. Adaptability within neuronal circuitry may influence learning and memory processes and may be important across the lifespan (Douglas Fields, 2015; Timmler & Simons, 2019).

Studies investigating the myelination response to activity-dependent plasticity have predominantly focused on early postnatal development and often used artificial neuronal stimulation paradigms (Gibson et al., 2014; Hill, Patel, Goncalves, Grutzendler, & Nishiyama, 2014; Makinodan et al., 2012; Mensch et al., 2015; Stanislaw Mitew et al., 2018). Optogenetic (Gibson et al., 2014) and pharmacogenetic (Stanislaw Mitew et al., 2018) methods to upregulate neuronal activity during postnatal development revealed greatly increased oligodendrocyte progenitor cell (OPC) proliferation and subsequent maturation, an affect which was recapitulated, albeit to a lesser extent, in young adult animals (Stanislaw Mitew et al., 2018). However, these artificial methods to upregulate neuronal stimulation to promote myelin plasticity may be problematic, as they may not actually reflect physiological levels of stimulation and have recently been shown to promote the progression of tumor growth (Venkataramani et al., 2019; Venkatesh et al., 2019). Interestingly, young adult mice (Xiao et al., 2016) and rats (Keiner et al., 2017) that were taught complex motor paradigms exhibited an increase in new OL production, suggesting that OL adaptability extends to adulthood and is physiologically relevant. While these findings are exciting, a complete understanding of the precise changes in cellular dynamics induced by physiologically relevant stimulation and how this ultimately influences myelin in the adult CNS remains unknown. Whether increases in OL production are driven by new OPC proliferation and subsequent differentiation, or the direct differentiation of pre-existing oligodendroglia, is yet to be comprehensively assessed. Furthermore, how the maintenance and structure of existing myelin sheaths and myelinated axons is altered and how the underlying cellular changes ultimately influence new myelin generation, is unknown. This warrants a deeper exploration of how physiologically relevant, activity-dependent myelination and oligodendroglial adaptations occur at post-developmental time points.

Environmental enrichment (EE) is a non-invasive and well-studied housing paradigm that is known to induce neuroplasticity and improve cognitive function (Nithianantharajah & Hannan, 2006). Using EE as a paradigm of physiologically relevant, activity-dependent plasticity, we investigated the generation of new myelin along with the ultrastructure of existing myelin sheaths and comprehensively assessed the production and differentiation of oligodendroglia, in 9-week-old mice. We found that 6 weeks of EE did not induce significant *de novo* generation of myelin sheaths on previously unmyelinated segments, but rather led to a generalized increase in axon diameter of already myelinated axons accompanied by thicker myelin sheaths in the corpus callosum, indicative of a remodeling of pre-existing myelinated axons. Further, EE uniformly accelerated the direct differentiation of predominantly pre-existing oligodendroglia in both the corpus callosum and somatosensory cortex, but differentially impeded OPC homeostasis in these regions. Together, our data indicate that physiologically relevant activity-dependent plasticity in the young adult CNS involves the remodeling of myelinated axons and promotes the direct differentiation of pre-existing oligodendroglia. Results of this study provide new insights into the dynamics of activity-dependent myelination in the adult CNS, opening up new questions concerning the life-long importance of myelinated axon remodeling and oligodendroglial differentiation in optimization and maintenance of CNS function.

## Materials and methods

### Experimental animals

C57BL/6 mice were bred and housed in specific pathogen-free conditions at the Melbourne Brain Centre Animal Facility, in Techniplast IVC cages (Techniplast Group, Italy). All procedures were approved by the Florey Institute for Neuroscience and Mental Health Animal Ethics Committee following, the Australian Code of Practice for the Care and Use of Animals for Scientific Purposes.

### Housing conditions and EdU administration

At 9 weeks of age, male and female animals were randomly assigned to standard or environmentally enriched housing conditions and housed single-sex with 3 mice/cage for a period of 6 weeks. Standard-housed mice remained in shoe-box sized GM500 cages (Techniplast Group, Italy) with floor area of 501cm^2^. Enriched mice were moved to GR1800 double-decker cages (Techniplast Group, Italy) with floor area of 1862cm^2^ and total height of 38cm. Enriched cages included a mouse-house and a selection of rubber dog toys, small plastic objects, tunnels and climbing materials. Standard-housed mice were provided only basic bedding materials and enriched mice had additional materials including shredded paper and cardboard. All cages were changed weekly with objects in enriched cages replaced, to maintain object novelty. All mice were given access to food and water ad libitum, and were on a 12-hour light/dark cycle.

Throughout the total housing period of 6 weeks, the drinking water contained thymidine analogue EdU (5-ethynyl-2’-deoxyuridine, Thermo Fisher, cat. no: E10415) to label newly-generated cells. EdU was at a concentration of 0.2mg/ml, determined previously to be non-toxic (Young et al., 2013), and refreshed every 2-3 days.

### Tissue collection

Mice were anesthetized and transcardially perfused with 0.1M phosphate buffer (PBS) followed by 4% EM-grade paraformaldehyde (PFA, Electron Microscopy Sciences). Brains were dissected and post-fixed overnight in 4% PFA. The first millimeter of caudal corpus callosum was selected using a coronal mouse brain matrix and micro-dissected, then placed in Kanovsky’s buffer overnight, washed in 0.1M sodium cacodylate and embedded in the sagittal plane in epoxy resin for transmission electron microscopy (TEM) analysis. The remaining brain was rinsed in 0.1M PBS, left overnight in a 30% sucrose solution to induce cryoprotection, then embedded in OCT and snap-frozen in isopentane over dry ice, for immunohistochemistry analysis.

### Immunohistochemistry and EdU labelling

20μm coronal cryosections at approximately Bregma −1.64mm were collected in series on SuperFrost plus slides. Sections were incubated overnight at room temperature with primary antibodies including, rabbit anti-Olig2 (Millipore, #AB9610, 1:200), mouse anti-APC/CC1 (CalBioChem, #OP-80, 1:200), and goat anti-platelet derived growth factor receptor-alpha (PDGFRα, R&D systems #AF1062, 1:200), in a PBS-based diluent buffer containing 10% normal donkey serum and 0.2% Triton-X100, then washed with 0.1M PBS and dark-incubated for 4h at room temperature with appropriate secondary antibodies including, donkey anti-rabbit 594 and donkey anti-goat 555 (Alexa Fluor, Invitrogen, 1:200) and donkey anti-mouse 405 (Abcam, ab175658, 1:200), in the same PBS-based diluent. Sections were washed with 0.1M PBS, and dark-incubated in the EdU developing cocktail prepared as per product instructions (Click-iT^™^ EdU Alexa Fluor^™^ 647 Imaging Kit, Thermo Fisher, cat. no: C10340) for 2h at room temperature, for EdU detection. Sections were washed a final time in 0.1M PBS and mounted in DAKO fluorescence mounting medium (Agilent Dako, cat. no: S3023).

### Spectral unmixing confocal microscopy imaging and cell counting

Images were obtained using a 20x objective on a Zeiss LSM880 laser scanning confocal microscope with 405nm, 561nm and 633nm laser lines and Zen Black 2.3 image acquisition software. Images were taken in Lamda mode, using a ChS1 spectral GaAsP (gallium arsenide phosphide) detector to capture the entire spectrum of light, generating one image containing all fluorophores. An individual spectrum for each fluorophore was obtained by imaging control, single-stained slides and these spectra used to segregate each multi-fluorophore image into a 4-channel image, in a post-processing step under the linear unmixing function. Uniform settings were used across experiments. Consistent regions of interest (ROI) were maintained across animals; mid-line corpus callosum and primary somatosensory cortex at approximately Bregma −1.64mm, and 3-4 images per ROI per animal were taken.

Images were manually counted by assessors blinded to housing conditions, using ImageJ/FIJI software. Oligodendroglia were defined as Olig2+, OPCs as Olig2+/ PDGFRα+ double-labelled cells, mature OLs as Olig2+/CC1+ double-labelled cells and intermediate oligodendroglia as Olig2+/CC1-/ PDGFRα-. The area of corpus callosum and somatosensory cortex was measured in each image and counts normalized.

### SCoRe microscopy imaging and analysis

20μm thick coronal sections were imaged on a Zeiss LSM880 laser scanning confocal microscope with a 40x water immersion objective using 488nm, 561nm and 633nm laser lines passed through the Acousto-Optical Tunable Filters 488-640 filter/splitter and a 20/80 partially reflective mirror. Compact myelin reflected light that was then collected using three photodetectors set to collect narrow bands of light around the laser wavelengths. Uniform settings were used across experiments. Images were acquired in tile scans of 5μm z-stacks, at a minimum z-depth of 5μm from the surface. ROIs were consistent with spectral images and 3-4 images per ROI per animal were taken. Image analysis was performed in ImageJ/FIJI. Maximum intensity z-projection images were applied a minimum threshold cut-off and the resulting area of positive pixels in each ROI measured.

### Transmission electron microscopy and analysis

At approximately Bregma −2.30mm semi-thin (0.5-1μm) sections of caudal corpus callosum were collected on glass slides in the sagittal plane and stained with 1% toluidine blue, for ROI identification. Subsequent ultrathin (0.1μm) sections were collected on 3×3mm copper grids and contrasted with heavy metals. Ultrathin sections were viewed using a JEOL JEM-1400Flash TEM. Images were captured using the JEOL integrated software and a high-sensitivity sCMOS camera (JEOL Matataki Flash). Eight-ten distinct fields of view were imaged at 10,000x magnification per animal. FIJI/Image J image analysis software (National Institutes of Health) was used to count myelinated axons and the Trainable WEKA Segmentation plugin (Leslie & Heese, 2017) used to segregate myelin, allowing use of the magic wand tool to measure inner and outer axon areas, to obtain axon diameter distribution and to calculate inner and outer diameters for g-ratio calculation. For each animal, at least 100 axons were measured. Resin embedding, sectioning, post-staining and EM imaging was performed at the Peter MacCallum Centre for Advanced Histology and Microscopy.

### Statistical analysis

All data was analyzed and graphed in GraphPad Prism vs8 software. Data were assumed to be normally distributed and variance assumed to be equal between groups. Sample size was determined based on what is generally used in the field. EM data was analyzed using a 2-way ANOVA with Sidak’s multiple comparison test or un-paired two-tailed t-test. Spectral and SCoRe imaging data was analyzed using an un-paired two-tailed t-test. Data are presented as mean ± SEM. A significance threshold *p*-value of 0.05 was used.

## Results

### Environmental enrichment increases axonal caliber and promotes growth of pre-existing myelin sheaths

To physiologically induce neuronal activity, we adopted the well-established EE housing paradigm (Nithianantharajah & Hannan, 2006). 9-week-old mice were housed in EE or standard-housed (SH) control conditions for a period of 6-weeks and administered the thymidine analogue EdU in the drinking water throughout to label newly-generated cells. To verify this housing paradigm, we confirmed neurogenesis in the hippocampus, a well-defined read out of EE (Nithianantharajah & Hannan, 2006). We observed a significant increase in the density of EdU+ cells in the dentate gyrus of EE mice compared to SH control mice (862 ± 108.7 cells/mm^2^ compared to 453 ± 10.1 cells/mm^2^, p=0.02, Student’s t-test, n=3-4 mice/group, data = mean ± SEM).

We first investigated the influence that EE exerts on myelin plasticity by assessing the myelinated area in the corpus callosum (Fig. 1A) using SCoRe microscopy, a reflection signal of compact myelin generated by incident lasers of multiple wavelengths (Gonsalvez et al., 2019; Hill et al., 2018; Schain, Hill, & Grutzendler, 2014). There was no change in the overall percentage area of myelin coverage between EE and SH control mice in the corpus callosum (Fig. 1A, quantitated in Fig. 1C), indicating that EE induced no significant increase in the *de novo* myelination of previously unmyelinated axons. Due to recent observations of continued myelin generation in the cortex throughout the lifespan (Hill et al., 2018) and acknowledgement that grey matter myelination is emerging as a key area of myelin plasticity (Timmler & Simons, 2019), we applied SCoRe microscopy imaging to the overlying region of the somatosensory cortex, from which the callosal projection fibers originate (Fig. 1B). Similar to the corpus callosum, we observed no significant change in the percentage area of myelin coverage following EE housing in this region (Fig. 1B, quantitated in Fig. 1D). Additional layer-specific analysis revealed that deeper cortical layers were more heavily myelinated than superficial layers (Fig. 1E, p=0.006), confirming the specificity of SCoRe as a myelin imaging technique, however no region-dependent difference was found between housing conditions. Collectively, these data suggest there is no significant increase in the *de novo* myelination of previously unmyelinated axons following EE in either corpus callosum or somatosensory cortex.

**Figure 1.**
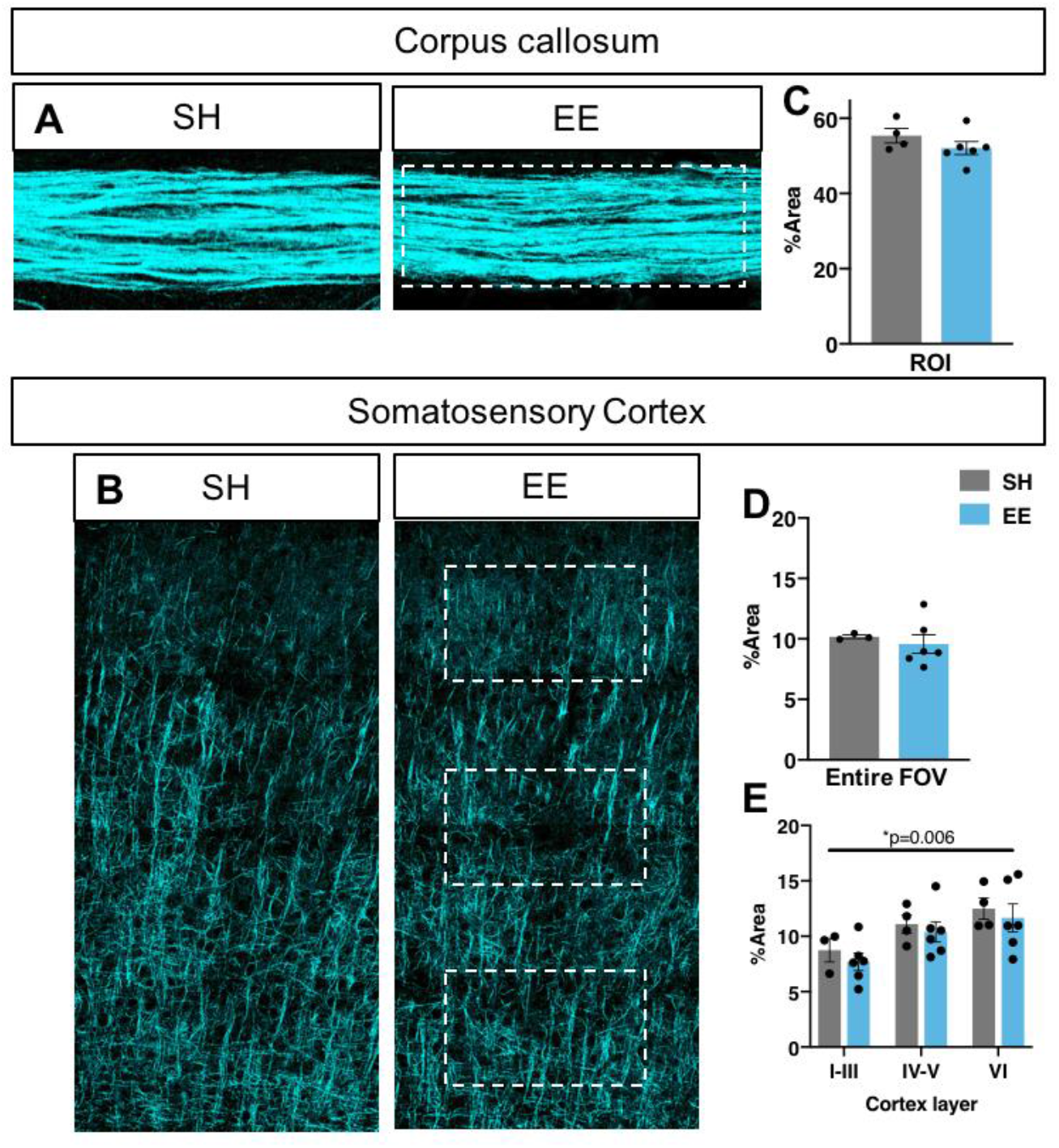
Environmental enrichment exerts no influence on percentage myelinated area. **(A-B)** Representative SCoRe images of corpus callosum **(A)** and somatosensory cortex **(B)** of control SH mice and EE housed mice. **(C-E)** Quantification of myelinated area as per percentage area of SCoRe signal measured in a corpus callosum ROI (p=0.25, unpaired two-tailed t-test) **(C)**, across the entire somatosensory cortex (p=0.62, unpaired two-tailed t-test) **(D)** and in 3 layer-specific cortical ROIs (dotted outlines, 2-way ANOVA with Sidak’s multiple comparisons) **(E)**. C-E: n=3-6 mice/group, data = mean ± SEM.

While SCoRe imaging can capture a gross increase of myelin production, the light reflective nature of this method for assessing myelin does not equip it to examine some important parameters of myelination such as individual sheath thickness and axon structure. Therefore, in order to more precisely investigate whether EE induces activity-dependent changes to myelin ensheathment, we assessed myelin ultrastructure using TEM in the corpus callosum (Fig. 2A). Concordant with the SCoRe imaging data, we found no significant change in the percentage of myelinated axons between housing conditions (Fig. 2B), confirming that EE exerted no influence on *de novo* myelination of previously unmyelinated axons. Interestingly, analysis of the frequency distribution of myelinated axons relative to their axonal diameter showed that EE mice had significantly larger axonal diameters compared to SH control (Fig. 2C, p=0.008). There was an approximately 30-50% reduction in the proportion of axons with small diameters (0.2-0.5μm) and a corresponding increase in the proportion of axons with large diameters (0.8-2.3μm), resulting in an overall ~30% increase in the mean diameter of myelinated axons in EE mice compared to SH control mice (Fig. 2D, p=0.02). More intriguingly, in addition to axonal caliber changes we also found a significant reduction in overall g-ratios in EE mice compared to SH controls (Fig. 2E, p<0.0001), indicative of thicker myelin sheaths. This reduction in g-ratios was observed across the full range of axon diameters (Fig. 2F, p<0.0001), indicating a generalized effect of EE in promoting the re-initiation of growth within existing myelin sheaths. Previous studies have identified an increased corpus callosum volume following EE housing (Markham, Herting, Luszpak, Juraska, & Greenough, 2009; Zhao et al., 2012), which we also observed via quantifying an increase in the vertical height of the corpus callosum in the coronal plane following EE (216.0 ± 6.07μm compared to 143.2 ± 1.30μm p<0.0001, Student’s t-test, n=3-4 mice/group, data = mean ± SEM). Our results provide clear evidence that young adult CNS myelination is highly adaptive to activity-dependent plasticity induced by enhanced environmental experiences. Importantly, we show this response does not involve an overt generation of *de novo* myelin sheaths on previously unmyelinated axon segments, but instead a remodeling of pre-existing myelinated axons, whereby axon diameters restructure and secondary growth of the accompanying myelin sheath is initiated.

**Figure 2.**
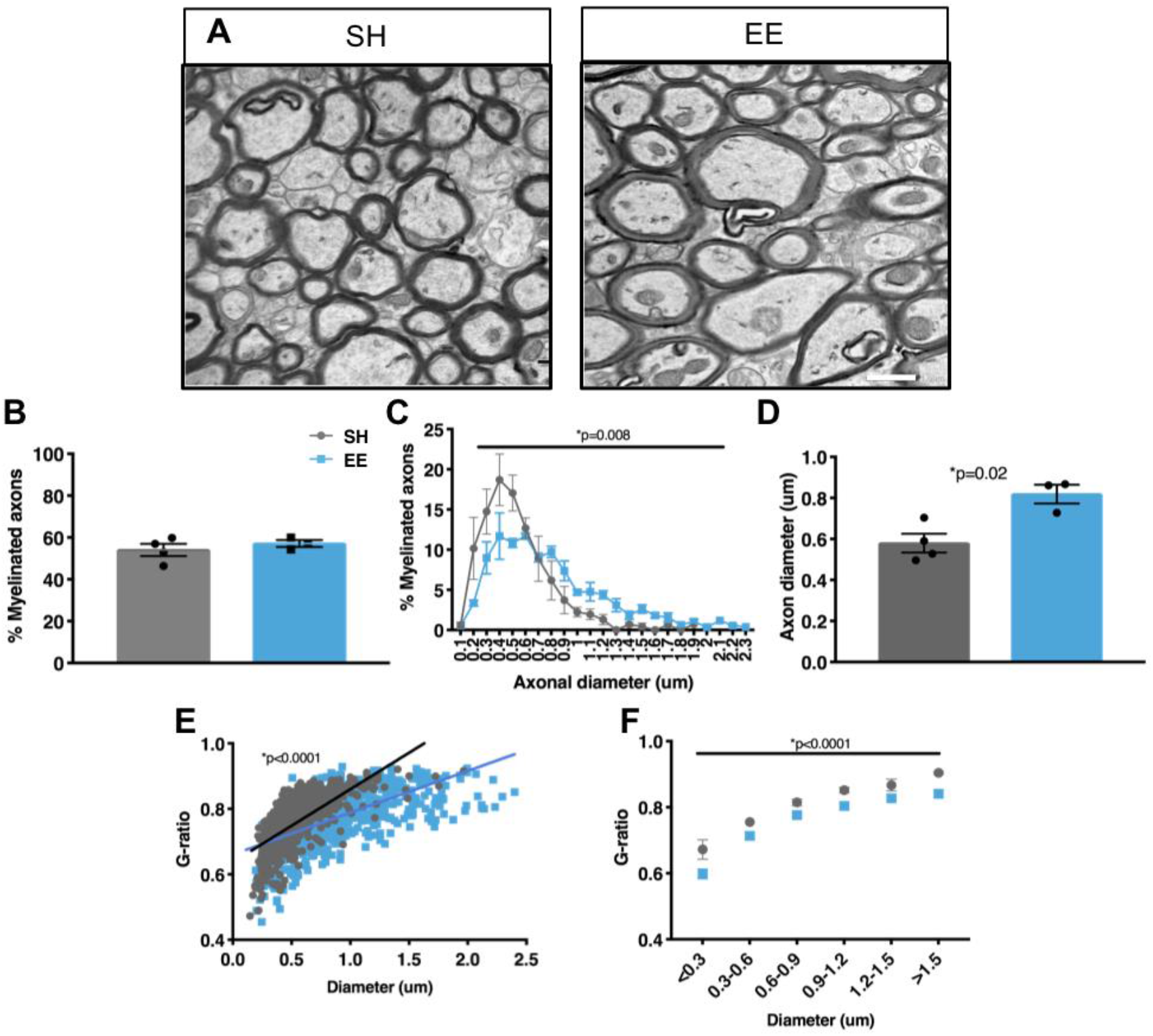
Environmental enrichment increases axonal caliber and thickness of pre-existing myelin sheaths in the corpus callosum. **(A)** Representative EM micrographs of caudal corpus callosum of control SH or EE housed mice. Scale bar = 1μm. **(B)** Quantification of the percentage of myelinated axons in the corpus callosum (p=0.45, unpaired two-tailed t-test). **(C)** Frequency distribution of myelinated axons relative to diameter (2-way ANOVA; interaction p=0.008). **(D)** Quantification of average axon diameter in the corpus callosum (p=0.02, unpaired two-tailed t-test). **(E)** Scatter plot of g-ratio distribution of individual axons relative to axon diameter (linear regression analysis, p<0.0001). **(F)** Plot of average g-ratio binned by axon diameter (2-way ANOVA with Sidak’s multiple comparison test; factors: housing p<0.0001, diameter p<0.0001, interaction p=0.8). For C-F: >100 axons/mouse/group, n=3-4 mice/group, data = mean+SEM.

### Environmental enrichment promotes the direct differentiation of pre-existing oligodendroglia in the corpus callosum and somatosensory cortex

To investigate the effect of EE on the cellular dynamics of OL production, we used spectral imaging confocal microscopy with linear unmixing and comprehensively analyzed the density and proportion of oligodendroglia. We assessed total oligodendroglia (Olig2+), OPCs (Olig2+/Pdgfrα+), intermediate oligodendroglia (Olig2+/Pdgfrα-/CC1-) and post-mitotic, mature OLs (Olig2+/CC1+), distinguishing between cells that were pre-existing (EdU-) and newly-generated (EdU+) during the 6-week EE period (Fig. 3A).

**Figure 3.**
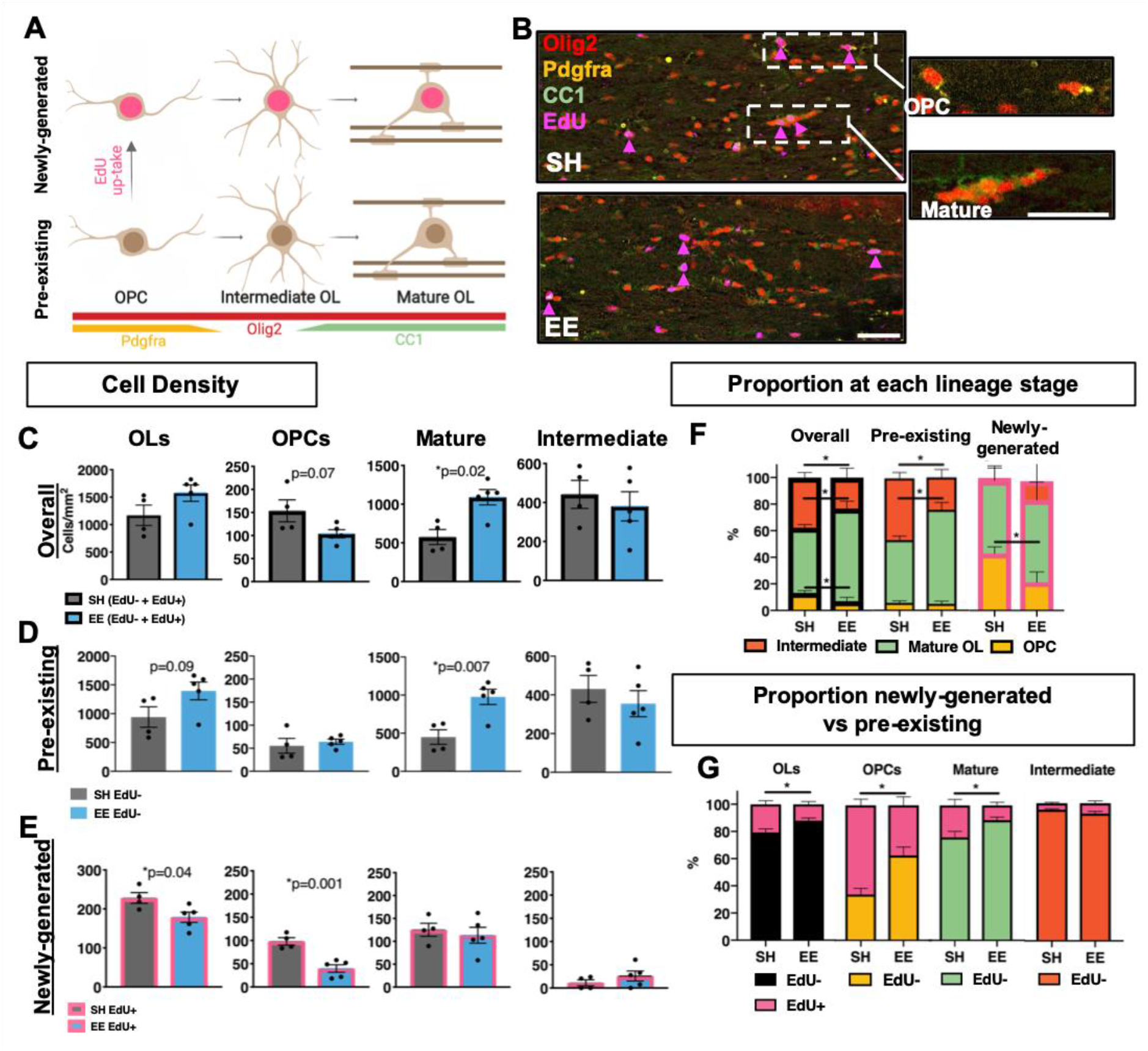
Environmental enrichment enhances the direct differentiation of pre-existing oligodendrocytes in the corpus callosum. **(A)** Schematic of the markers used to label distinct stages of the oligodendroglial lineage. This image was created using BioRender.com. **(B)** Representative confocal micrographs of Olig2/EdU/PDGFRAα/CC1 quadruple-immunostaining in the corpus callosum for SH or EE housed mice. Insert from SH animal depicts new-OLs (EdU+, pink arrows), OPCs (Olig2+/PDGFRAα+) and mature OLs (Olig2+/CC1+). Scale bar = 25μm. **(C-E)** Quantification of the density of total oligodendroglia (Olig2+), as well as oligodendroglia sub-divided by lineage stage; OPCs (Olig2+/ PDGFRAα+), mature OLs (Olig2+/CC1+) and intermediate oligodendroglia (Olig2+/ PDGFRAα-/CC1-) in the corpus callosum. **(C)** Overall oligodendroglial densities [Pre-existing (EdU-) + Newly-generated (EdU+)] (OLs p=0.13, OPCs p=0.07, Mature p=0.02, Intermediate p=0.58). **(D)** Pre-existing (EdU-) oligodendroglial densities (OLs p=0.09, OPCs p=0.58, Mature p=0.007, Intermediate p=0.46). **(E)** Newly-generated (EdU+) oligodendroglial densities (OLs p=0.04, OPCs p=0.001, Mature p=0.62, Intermediate p=0.21). **(F)** Quantification of the proportions of oligodendroglia at each stage of lineage development, for the overall population of oligodendroglia (OPCs p=0.006, Mature p=0.0002, Intermediates p=0.008), the population of pre-existing (EdU-) oligodendroglia (OPCs p=0.57, Mature p=0.0001, Intermediates p=0.0005) and the population of newly-generated (EdU+) oligodendroglia (OPCs p=0.002, Mature p=0.39, Intermediate p=0.18). **(G)** Quantification of the proportions of newly-generated (EdU+) vs pre-existing (EdU-) oligodendroglia overall (p=0.03), and at each lineage stage in the corpus callosum (OPCs p=0.009, Mature p=0.03, Intermediate p=0.30). C-G: n=4-5 mice/group, unpaired two-tailed t-test, data = mean + SEM.

To align with our myelin data, identical, adjacent regions of corpus callosum and somatosensory cortex were analyzed. In both regions, we observed a clear increase in the density of mature OLs (callosum: Fig. 3C, p=0.02; cortex: Fig. 4B p=0.02) in EE mice compared to SH controls, indicative of increased oligodendrocyte differentiation. This effect was more pronounced in the cortex, where EE led to an increase in overall OL density (Fig. 4B, p=0.04) compared to SH controls. This finding is concordant with previous observations of increased differentiation following activity-dependent stimulation (Cullen et al., 2019; Gibson et al., 2014; Hughes, Orthmann-Murphy, Langseth, & Bergles, 2018; Keiner et al., 2017; Stanislaw Mitew et al., 2018; Xiao et al., 2016). To expand on these studies, we wanted to precisely define the cellular dynamics underlying increased mature OL production induced by EE, and segregated the densities of all oligodendrocytes into both pre-existing (EdU-) and newly-generated (EdU+) populations (callosum: Fig. 3D-E; cortex: Fig. 4C-D). This approach was used to deduce which oligodendroglial population was primarily responsive to physiological EE stimulation.

**Figure 4.**
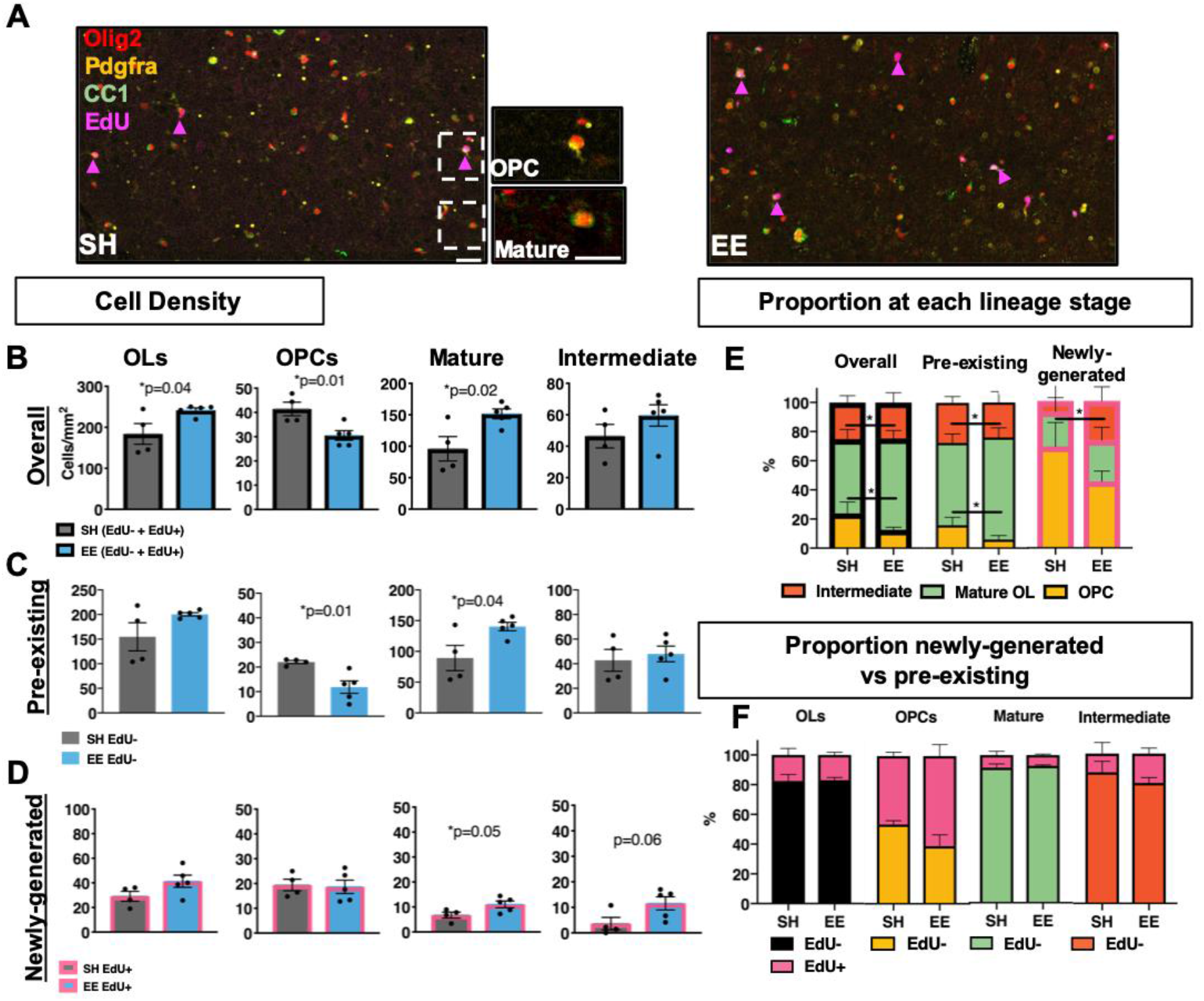
Environmental enrichment enhances the direct differentiation of pre-existing and new oligodendroglia in the somatosensory cortex. **(A)** Representative confocal micrographs of Olig2/EdU/PDGFRAα/CC1 quadruple-immunostaining in the somatosensory cortex for SH or EE house mice. Insert from SH animal depicts new-oligodendroglia (EdU+, pink arrows), OPCs (Olig2+/PDGFRAα+) and mature OLs (Olig2+/CC1+). Scale bar = 25μm. **(B-D)** Quantification of the density of total oligodendroglia, as well as oligodendroglia sub-divided by lineage stage; OPCs, mature OLs and intermediate oligodendroglia in the somatosensory cortex. **(B)** Overall oligodendroglial densities (OLs p=0.04, OPCs p=0.01, Mature p=0.02, Intermediate p=0.24). **(C)** Pre-existing oligodendroglial densities (OLs p=0.11, OPCs p=0.01, Mature p=0.04, Intermediate p=0.64). **(D)** Newly-generated oligodendroglial densities (OLs p=0.11, OPCs p=0.85, Mature p=0.05, Intermediate p=0.06). **(E)** Quantification of the proportions of oligodendroglia at each stage of lineage development, for the overall population of oligodendroglia (OPCs: p=0.02, Mature: p=0.02, Intermediate p=0.9), the population of pre-existing (EdU-) oligodendroglia (OPCs: p=0.01, Mature: p=0.02, Intermediate p=0.44) and the population of newly-generated (EdU+) oligodendroglia (OPCs: p=0.02, Mature p=0.51, Intermediate p=0.07). **(F)** Quantification of the proportions of newly-generated (EdU+) vs pre-existing (EdU-) oligodendroglia overall (p=0.91) and at each lineage stage in the somatosensory cortex (OPCs p=0.15, Mature p=0.60, Intermediate p=0.36). B-F: n=4-5 mice/group, unpaired two-tailed t-test, data = mean ± SEM.

We found an increased density of pre-existing mature OLs in EE mice compared to SH controls in both regions (callosum: Fig. 3D, p=0.007; cortex: Fig. 4C p=0.04), suggesting that EE uniformly increases the direct differentiation of pre-existing oligodendroglia. We also analyzed the proportions of sub-populations of oligodendroglia at each lineage phase (callosum: Fig. 3F; cortex: Fig. 4E) to further ascertain the increased maturation. Intuitively, our data revealed an increase in the proportion of mature OLs in both regions within both the total overall oligodendroglial cell pool (callosum: Fig. 3F, p=0.0002; cortex: Fig 4E, p=0.02) and specifically in the pre-existing oligodendroglial population (callosum: Fig. 3F, p=0.0001; cortex: Fig 4E, p=0.02), indicative of a robust differentiation. Collectively, our data indicate that the young adult brain contains an accessible reserve of oligodendroglia for activity-dependent responsiveness and that the lineage progression of pre-existing cells is particularly relevant for activity-dependent myelin plasticity induced by physiological stimuli in the young adult brain.

### Environmental enrichment alters OPC homeostasis is in the corpus callosum and somatosensory cortex

Having identified that EE uniformly increases the density of mature OLs in both corpus callosum and somatosensory cortex via enhancing differentiation of pre-existing cells, we found an associated effect on OPC homeostasis with a uniform decrease in the density of total OPCs in EE mice compared to SH controls. Interestingly, in the corpus callosum this decrease is specific to the density of newly-generated OPCs (Fig. 3E, p=0.001), whereas in the somatosensory cortex, this reduction is specific to pre-existing OPCs (Fig. 4C, p=0.01). There is evidence of increased differentiation within this population of newly-generated cells in response to EE, as per an increase in density of newly-generated mature OLs in the somatosensory cortex (Fig. 4D, p=0.05) and a decrease in proportion of OPCs in both regions (callosum: Fig. 3F, p=0.002; cortex: Fig. 4E, p=0.02).

To further determine the regionally-distinct reduction in new oligodendrocyte generation we examined the proportional contributions of new vs pre-existing oligodendroglia to the pool of total oligodendroglia, OPCs, mature OLs and intermediate oligodendroglia in both regions. In the corpus callosum, the overall proportion of pre-existing oligodendroglia was significantly increased in EE mice compared to SH controls (Fig. 3G, p=0.03), which is reflected by increased proportions of both pre-existing OPCs (Fig. 3G, p=0.009) and pre-existing mature OLs (Fig. 3G, p=0.03). Interestingly, EE exerted no significant influence upon the relative proportions of pre-existing and new oligodendroglia in somatosensory cortex (Fig. 4F), indicating clear regional heterogeneity. Together, our data indicate that prolonged, physiological stimulation via EE exerts a unified effect upon potentiating the differentiation of pre-existing oligodendroglia in both the corpus callosum and somatosensory cortex and differentially alters the OPC homeostasis between the two regions.

## Discussion

This study reports a previously undescribed form of activity-dependent adaptive myelination in the young adult CNS. Results of this study identify that physiological stimulation via EE exerts little effect on the proliferation of new oligodendrocytes or the *de novo* generation of new myelin sheaths on previously unmyelinated axons, but increases the direct differentiation of pre-existing oligodendrocytes and induces the remodeling of axon-myelin units via enlargement of axon caliber and secondary growth of the overlying, pre-existing myelin sheaths. Together, these findings illustrate a different form of activity-dependent plasticity within oligodendroglia and CNS myelin in response to physiological stimuli, raising mechanistic questions for future investigations.

In this current study, we did not observe significant *de novo* generation of myelin sheaths upon previously unmyelinated segments, a finding which is perhaps unsurprising, given that by P90 the proportion of myelinated axons in the murine corpus callosum has almost peaked (Sturrock, 1980). Although generation of *de novo* myelination continues throughout the lifespan, particularly in the cortex (Hill et al., 2018) and it could be assumed that each new oligodendrocyte produces it’s average of 50 new sheaths (Hughes et al., 2018), the relative contribution of myelin generated over this period is marginal, perhaps rendering bulk analysis of total myelinated area insensitive. The activity-dependent myelination field currently focuses heavily on the *de novo* myelination of previously unmyelinated axons and has not yet thoroughly considered the possibility that myelin plasticity, particularly in adult animals, involves subtle, structural modulation of existing myelin sheaths, such as thickness changes (Lazari, Koudelka, & Sampaio-Baptista, 2018). This is most likely due to technical limitations with no current common techniques enabling simultaneous analysis of myelin sheath length and myelin thickness, however the recent surge in cryo-EM may provide a powerful solution. If used in combination with a genetically encoded, inducible, membrane-bound, fluorescent reporter, this would enable specific comparison between *de novo* generated sheaths and pre-existing sheaths, to definitively answer this question.

Most interestingly in this current study, we provide initial evidence of such axon-myelin unit plasticity and we are the first to do so in a physiologically induced, *in vivo* context. We demonstrate a significant and generalized effect of axon diameter enlargement, accompanied by an increase in thickness of the overlying myelin sheaths. High-frequency stimulation of hippocampal neurons has been recently shown to enlarge synaptic boutons and correspondingly induce an increase in axon diameters by ~5% upon a time scale of 10-30 minutes (Chéreau, Saraceno, Angibaud, Cattaert, & Nägerl, 2017). Although perhaps mechanistically different, our data further confirm structural plasticity within axons and suggest that over a sustained period of physiologically relevant stimulation, the overall axon diameter can continue to increase up to 30%. Activity-dependent axonal structural changes have been studied very little (Costa, Pinto-Costa, Sousa, & Sousa, 2018; Lazari et al., 2018) and to our knowledge, we are the first to show a long-term, physiologically-induced axon caliber growth *in vivo*. In conjunction with axon caliber enlargement we found thicker myelin as result of an apparent re-initiation of secondary myelin growth within the surrounding, pre-existing sheaths. Increasing evidence suggests that myelin remains a dynamic structure throughout the lifespan, as observed by fluctuations in internode length (Hill et al., 2018) and turnover of myelin (Lüders et al., 2019; Yeung et al., 2014), implying there are molecular substrates susceptible to adaptive regulation. A recent study has demonstrated activity-dependent localization of myelin protein mbpa mRNA within developing myelin sheaths (Yergert, Hines, & Appel, 2019), indicating a potential molecular correlate of localized sheath growth. We know that myelin growth during development involves the inner tongue enlarging and growing under the overlying layer (Snaidero & Simons, 2017), implying existing logistics to support secondary myelin growth and in contexts of exogenous stimulation, specific activity-dependent increases in myelin sheath thickness has been shown after optogenetic (Gibson et al., 2014) and chemogenetic DREADD-mediated (Stanislaw Mitew et al., 2018) stimulation. Plasticity inherently implies bidirectionality and indeed myelin thinning has been observed following social isolation (Liu et al., 2012; Makinodan et al., 2012), chronic social stress (Bonnefil et al., 2019) and loss of auditory stimulation (Sinclair et al., 2017). Secondary myelin growth has also been initiated via genetic manipulation involving conditional deletion of Pten and subsequent elevation of PI(3,4,5)P3 in adult (P60) optic nerve (Snaidero et al., 2014), interestingly a mechanism also known to increase axonal caliber (Goebbels et al., 2016). It could easily be speculated that this activity-dependent sheath growth is in fact inherently linked to axon caliber growth, as although not quantified, in representative images it appears that stimulated axons have larger diameters (S. Mitew et al., 2014) and loss of auditory stimulation was in fact associated with a reduction in the number of large diameter fibers (Sinclair et al., 2017), suggesting an intrinsically interlinked mechanism of both axon caliber and myelin sheath growth.

Indeed, it is interesting to speculate on the interconnected mechanism and potential functional consequences of the concurrent increase in axon diameter and myelin sheath thickness. Axon caliber is dictated by the axonal cytoskeleton, which functionally regulates the essential antero-and retrograde transport (Leterrier, Dubey, & Roy, 2017) and axon caliber is known to be dynamic, with structural mechanisms likely involving cytoskeletal reorganization, reviewed (Costa et al., 2018). The essentiality of cytoskeletal stability for axonal transport and synaptic function was demonstrated by work in *Drosophila* (Stephan et al., 2015) and it is known that a break-down in axonal transport is a pathological feature of a whole host of adult-onset neurodegenerative diseases (Brady & Morfini, 2017). Developmentally, local regulation between axon and sheath enable a single oligodendrocyte to ensheath axons of various caliber to the appropriate thickness (Waxman & Sims, 1984). Furthermore, the expansion of axon diameter may in fact be reliant on myelin, as during retinal ganglion development oligodendrocyte signals trigger local neurofilament accumulation and network reorganization to enable axon enlargement (Sánchez, Hassinger, Paskevich, Shine, & Nixon, 1996) and unhealthy myelin induced by sulfatide loss leads to axonal caliber reduction and break down (Marcus et al., 2006), suggesting the thickness and integrity of myelin is critical in maintaining larger axonal diameters. There is yet to be definitive evidence for linking oligodendrocytes and myelination to axonal caliber maintenance and axonal transport function, however it is known that oligodendrocytes secrete extracellular vesicles and these are known to enhance neuronal physiology via increasing stress tolerance and firing rate (Fröhlich & Kuo, 2014). These vesicles have recently been shown to facilitate axonal transport and myelin proteins PLP- and CNP-deficient mice have deficiencies in this process, coupled to secondary axonal degeneration, providing a potential mechanism linking myelin dysfunction to axonal transport impairment and thus maintenance of optimal neuronal function (Frühbeis et al., 2019). There is, however, a well-established role for oligodendrocytes and myelin in providing metabolic support to neurons by supplying pyruvate and lactate (Fünfschilling et al., 2012; Lee et al., 2012) and this is known to be regulated in an activity-dependent manner (Saab et al., 2016). As activity rates and axon caliber are linked (Perge, Niven, Mugnaini, Balasubramanian, & Sterling, 2012), it is highly likely that our findings of increased myelin thickness have implications for the amount of metabolic support provided to underlying, enlarged neurons and indeed emerging evidence suggests this is necessary in sustaining function of highly active circuits (Moore et al., 2019).

In this current study we performed a comprehensive analysis of oligodendroglial lineage progression and observed primarily that the physiological oligodendroglial response to activity-dependent plasticity involves the direct differentiation of pre-existing cells. This cellular response is likely explained by an activity-induced decrease in homeostatic cellular death, as it has been shown previously that EE enhances the survival of newly-generated oligodendroglia in the amygdala (Okuda et al., 2009). Indeed, elegant classification of homeostatic oligodendroglial differentiation has previously demonstrated that a continual turnover, involving loss of and subsequent proliferation, of OPCs occurs in the adult somatosensory cortex (Hughes, Kang, Fukaya, & Bergles, 2013). The basal differentiation and integration of OPCs has been quantified at merely 22%, but with sensory stimulation can be increased 5-fold (Hughes et al., 2018), suggesting activity-dependent mechanisms of retention of these cells. Similarly, stimulating neuronal circuits via transcranial magnetic stimulation results in more new pre-myelinating OLs in the motor and visual cortices (Cullen et al., 2019) and stimulating via complex motor learning enhances the differentiation and integration of pre-existing cells (captured via EdU administration prior to paradigm onset) (Xiao et al., 2016). In conjunction with published work, this study suggests that pre-myelinating oligodendroglia form an accessible reserve of responsive cells in the young adult brain that are amenable to drive adaptive regulation. The continued differentiation and survival of these cells appears to be key for physiological myelin plasticity in adulthood, which stands contrary to the evidence collected previously in juvenile animals under artificial stimulation paradigms, which primarily reported activity-dependent responses involving the mass induction of OPC proliferation preceding subsequent differentiation.

Most curious in this current study is that in both regions of analysis there appears to be a reduction in the homeostatic proliferation of OPCs, as per reduction in density. Previous live-imaging characterization of OPC homeostasis revealed that the majority of OPCs directly differentiate without prior proliferation and that OPC density is maintained by neighboring OPC proliferation and replacement (Hughes et al., 2013), therefore it is puzzling as to why density was not thusly maintained in this situation. As EE is a paradigm of non-specific stimulation, it is possible that only sub-populations of neurons within our analysis regions are responsive to this stimulation, and it is a localized group of OPCs that are responding, with distal OPCs unchanged. However, we analyzed oligodendroglia at one final time point providing no indication of the temporal change in underlying cellular dynamic alterations and perhaps if returned to standard housing, homeostasis would return. It is interesting to speculate though, as our EE paradigm is relatively mild stimulation but experienced continually for a rather long, 6-week period, if it may be that slowly the OPC pool is depleted, through a prolonged enhancement of differentiation that eventually begins to outpace replacement. This would indicate a physiological maximum of possible oligodendroglial responsiveness to activity in adulthood, which would have implications for designing remyelinating or other oligodendroglial-targeted therapies for adult-based brain injury or disease contexts. In fact, previous studies have identified that young adult animals (P60) (Mckenzie et al., 2014; Stanislaw Mitew et al., 2018) exhibit attenuated OPC production responses to neuronal stimulation as compared to juvenile animals (P19-35) (Gibson et al., 2014; Stanislaw Mitew et al., 2018), suggesting that age is a critical factor in the capacity of OPCs to produce large quantities of new cells. We have previously shown that there is a drastic decline in OPC density occurring across different CNS regions from P9-30 (Nicholson et al., 2018) and perhaps it is not that adult animals lack the ability to respond to neuronal activity, but rather that there are logistical constraints on cell numbers available to respond. Inherently, production and differentiation cannot occur simultaneously, as differentiation removes an available proliferative cell, and as a very tightly regulated cellular process it is difficult to imagine that proliferation or cell cycle length could be drastically shortened to more quickly produce greatly increased numbers of cells. In this study, we did observe some regional heterogeneity in OPC homeostasis and it is known that cortical and callosal OPCs behave differently from early postnatal development (Nicholson et al., 2018), with regional differences in subsets of OPCs having been identified based on differential expression of ion channels and neurotransmitter receptors (Spitzer et al., 2019). However in this context, heterogeneity may merely reflect the developmental timeline of myelination in each region, as over half of the total cortical oligodendrocytes are generated after 4 months of age (Hughes et al., 2018), a time well after the corpus callosum has reached peak myelination (Sturrock, 1980). Of note is that at 15 weeks of age, there are proportionally less OPCs in the control corpus callosum vs cortex, ~20% vs 10% (Fig. 3F and Fig. 4E), perhaps reflecting the later developmental stage in the corpus callosum vs cortex, with respect to level of myelination, and implying a greater logistical constraint on the extent of proliferation possible in the corpus callosum.

In summary, we for the first time demonstrate that in the young adult brain, activity-dependent myelin plasticity at a physiological level is predominantly driven by the remodeling of pre-existing myelinated axons and enhanced differentiation of pre-existing oligodendroglia, without overt *de novo* myelin sheath generation or mass OPC proliferation. These findings are important as they suggest that the pronounced *de novo* oligodendrogenesis as described by others may be more the result of non-physiological or supra-physiological stimuli, and the mechanisms underlying the remodeling of pre-existing myelinated axons and oligodendrocytes, along with the functional consequences thereof clearly warrant future investigations. Techniques to capture such specific changes could include methods that measure myelin generation and turnover, which were recently nicely reviewed (Buscham, Eichel, Siems, & Werner, 2019). Additionally, genetic approaches that enable inducible and OL-specific expression of a myelin-associated membrane-bound fluorescent protein (Hill & Grutzendler, 2019) in conjunction with time-lapse *in vivo* imaging, would provide a more detailed spatial and temporal analysis of the adaptive myelinating process. Together, results of this study add another layer of complexity to activity-dependent myelin plasticity and will provide new insights into the importance of the coincident physiological modulation of the axon-myelin unit in the adult CNS. Findings of this study will provide a platform for future investigation into the functional importance of myelinated axon plasticity for maintenance of optimum cognitive function across the lifespan.

## Acknowledgements

This study was supported by Australian Research Council Discovery Project Grant (#DP180102397) to Xiao J.; the Australian Government Research Training Program Scholarship and the University of Melbourne STRAPA Scholarship to M.N. Confocal Microscopy was performed at the Biological Optical Microscopy Platform, The University of Melbourne (www.microscopy.unimelb.edu.au).

## Notes

**Conflict of interest statement** The authors declare no conflicting financial or other interests.

### Competing Interest Statement

The authors have declared no competing interest.

## References

Bonnefil, V., Dietz, K., Amatruda, M., Wentling, M., Aubry, A. V, Dupree, J. L., … Liu, J. (2019). Region-specific myelin differences define behavioral consequences of chronic social defeat stress in mice. ELife, 8, 1–13. https://doi.org/10.7554/eLife.40855

Brady, S. T., & Morfini, G. A. (2017). Regulation of motor proteins, axonal transport deficits and adult-onset neurodegenerative diseases. Neurobiology of Disease, 105, 273–282. https://doi.org/10.1016/j.nbd.2017.04.010

Buscham, T., Eichel, M., Siems, S., & Werner, H. (2019). Turning to myelin turnover. Neural Regeneration Research, 14(12), 2063. https://doi.org/10.4103/1673-5374.262569

Castelfranco, A. M., & Hartline, D. K. (2015). The evolution of vertebrate and invertebrate myelin: a theoretical computational study. Journal of Computational Neuroscience, 38(3), 521–538. https://doi.org/10.1007/s10827-015-0552-x

Chéreau, R., Saraceno, G. E., Angibaud, J., Cattaert, D., & Nägerl, U. V. (2017). Superresolution imaging reveals activity-dependent plasticity of axon morphology linked to changes in action potential conduction velocity. Proceedings of the National Academy of Sciences of the United States of America, 114(6), 1401–1406. https://doi.org/10.1073/pnas.1607541114

Costa, A. R., Pinto-Costa, R., Sousa, S. C., & Sousa, M. M. (2018). The Regulation of Axon Diameter: From Axonal Circumferential Contractility to Activity-Dependent Axon Swelling. Frontiers in Molecular Neuroscience, 11(September), 1–7. https://doi.org/10.3389/fnmol.2018.00319

Cullen, C. L., Senesi, M., Tang, A. D., Clutterbuck, M. T., Auderset, L., O’Rourke, M. E., … Young, K. M. (2019). Low-intensity transcranial magnetic stimulation promotes the survival and maturation of newborn oligodendrocytes in the adult mouse brain. Glia, (March), glia.23620. https://doi.org/10.1002/glia.23620

Douglas Fields, R. (2015). A new mechanism of nervous system plasticity: activity-dependent myelination. Nature Reviews. Neuroscience, 16(12), 756–767. https://doi.org/10.1038/nrn4023

Foster, A. Y., Bujalka, H., & Emery, B. (2019). Axoglial interactions in myelin plasticity: Evaluating the relationship between neuronal activity and oligodendrocyte dynamics. Glia, (February), glia.23629. https://doi.org/10.1002/glia.23629

Fröhlich, D., & Kuo, W. D. (2014). Multifaceted effects of oligodendroglial exosomes on neurons. Retrieved from https://www.researchgate.net/profile/Eva-Maria_Kraemer-Albers/publication/264809938_Multifaceted_effects_of_oligodendroglial_exosomes_on_neurons_impact_on_neuronal_firing_rate_signal_transduction_and_gene_regulation/links/552ad6390cf2e089a3aa10bd.pdf

Frühbeis, C., Kuo-Elsner, W. P., Barth, K., Peris, L., Tenzer, S., Möbius, W., … Krämer-Albers, E.-M. (2019). Oligodendrocyte-derived exosomes promote axonal transport and axonal long-term maintenance. https://doi.org/10.1101/2019.12.20.884171

Fünfschilling, U., Supplie, L. M., Mahad, D., Boretius, S., Saab, A. S., Edgar, J., … Nave, K. A. (2012). Glycolytic oligodendrocytes maintain myelin and long-term axonal integrity. Nature, 485(7399), 517–521. https://doi.org/10.1038/nature11007

Gibson, E. M., Purger, D., Mount, C. W., Goldstein, A. K., Lin, G. L., Wood, L. S., … Monje, M. (2014). Neuronal Activity Promotes Oligodendrogenesis and Adaptive Myelination in the Mammalian Brain. Science, 344(6183), 1252304–1252304. https://doi.org/10.1126/science.1252304

Goebbels, S., Wieser, G. L., Pieper, A., Spitzer, S., Weege, B., Yan, K., … Nave, K.-A. (2016). A neuronal PI(3,4,5)P3-dependent program of oligodendrocyte precursor recruitment and myelination. Nature Neuroscience, 20(1), 10–15. https://doi.org/10.1038/nn.4425

Gonsalvez, D. G., Yoo, S. W., Fletcher, J. L., Wood, R. J., Craig, G. A., Murray, S. S., & Xiao, J. (2019). Imaging and Quantification of Myelin Integrity After Injury With Spectral Confocal Reflectance Microscopy. Frontiers in Molecular Neuroscience, 12(November), 1–13. https://doi.org/10.3389/fnmol.2019.00275

Hill, R. A., & Grutzendler, J. (2019). Uncovering the biology of myelin with optical imaging of the live brain. Glia, (February), glia.23635. https://doi.org/10.1002/glia.23635

Hill, R. A., Li, A. M., & Grutzendler, J. (2018). Lifelong cortical myelin plasticity and age-related degeneration in the live mammalian brain. Nature Neuroscience 2018, 26, 1. https://doi.org/10.1038/s41593-018-0120-6

Hill, R. A., Patel, K. D., Goncalves, C. M., Grutzendler, J., & Nishiyama, A. (2014). Modulation of oligodendrocyte generation during a critical temporal window after NG2 cell division. Nature Neuroscience, 17(11), 1518–1527. https://doi.org/10.1038/nn.3815

Hughes, E. G., Kang, S. H., Fukaya, M., & Bergles, D. E. (2013). Oligodendrocyte progenitors balance growth with self-repulsion to achieve homeostasis in the adult brain. Nature Neuroscience, 16(6), 668–676. https://doi.org/10.1038/nn.3390

Hughes, E. G., Orthmann-Murphy, J. L., Langseth, A. J., & Bergles, D. E. (2018). Myelin remodeling through experience-dependent oligodendrogenesis in the adult somatosensory cortex. Nature Neuroscience 2018, 1. https://doi.org/10.1038/s41593-018-0121-5

Keiner, S., Niv, F., Neumann, S., Steinbach, T., Schmeer, C., Hornung, K., … Redecker, C. (2017). Effect of skilled reaching training and enriched environment on generation of oligodendrocytes in the adult sensorimotor cortex and corpus callosum. BMC Neuroscience, 18(1), 31. https://doi.org/10.1186/s12868-017-0347-2

Lazari, A., Koudelka, S., & Sampaio-Baptista, C. (2018). Experience-related reductions of myelin and axon diameter in adulthood. Journal of Neurophysiology, 120(4), 1772–1775. https://doi.org/10.1152/jn.00070.2018

Lee, Y., Morrison, B. M., Li, Y., Lengacher, S., Farah, M. H., Hoffman, P. N., … Rothstein, J. D. (2012). Oligodendroglia metabolically support axons and contribute to neurodegeneration. Nature, 487(7408), 443–448. https://doi.org/10.1038/nature11314

Leslie, M. E., & Heese, A. (2017). Quantitative Analysis of Ligand-Induced Endocytosis of FLAGELLIN-SENSING 2 Using Automated Image Segmentation. In Trends in Immunology (Vol. 35, pp. 39–54). https://doi.org/10.1007/978-1-4939-6859-6_4

Leterrier, C., Dubey, P., & Roy, S. (2017). The nano-architecture of the axonal cytoskeleton. Nature Reviews Neuroscience, 18(12), 713–726. https://doi.org/10.1038/nrn.2017.129

Liu, J., Dietz, K., Deloyht, J. M., Pedre, X., Kelkar, D., Kaur, J., … Casaccia, P. (2012). Impaired adult myelination in the prefrontal cortex of socially isolated mice. Nature Neuroscience, 15(12), 1621–1623. https://doi.org/10.1038/nn.3263

Lüders, K. A., Nessler, S., Kusch, K., Patzig, J., Jung, R. B., Möbius, W., … Werner, H. B. (2019). Maintenance of high proteolipid protein level in adult central nervous system myelin is required to preserve the integrity of myelin and axons. Glia, 67(4), 634–649. https://doi.org/10.1002/glia.23549

Makinodan, M., Rosen, K. M., Ito, S., & Corfas, G. (2012). A Critical Period for Social Experience-Dependent Oligodendrocyte Maturation and Myelination. Science, 337(6100), 1357–1360. https://doi.org/10.1126/science.1220845

Marcus, J., Honigbaum, S., Shroff, S., Honke, K., Rosenbluth, J., & Dupree, J. L. (2006). Sulfatide is essential for the maintenance of CNS myelin and axon structure. Glia, 53(4), 372–381. https://doi.org/10.1002/glia.20292

Markham, J. A., Herting, M. M., Luszpak, A. E., Juraska, J. M., & Greenough, W. T. (2009). Myelination of the corpus callosum in male and female rats following complex environment housing during adulthood. Brain Research, 1288, 9–17. https://doi.org/10.1016/j.brainres.2009.06.087

Mckenzie, I. A., Ohayon, D., Li, H., Faria, J. P. De, Emery, B., Tohyama, K., & Richardson, W. D. (2014). Motor skill learning requires active central myelination. Science, 346(6207), 318–322. https://doi.org/10.1126/science.1254960

Mensch, S., Baraban, M., Almeida, R., Czopka, T., Ausborn, J., El Manira, A., & Lyons, D. A. (2015). Synaptic vesicle release regulates myelin sheath number of individual oligodendrocytes in vivo. Nature Neuroscience, 18(5), 628–630. https://doi.org/10.1038/nn.3991

Mitew, S., Hay, C. M., Peckham, H., Xiao, J., Koenning, M., & Emery, B. (2014). Mechanisms regulating the development of oligodendrocytes and central nervous system myelin. Neuroscience, 276(2014), 29–47. https://doi.org/10.1016/j.neuroscience.2013.11.029

Mitew, Stanislaw, Gobius, I., Fenlon, L. R., McDougall, S. J., Hawkes, D., Xing, Y. L., … Emery, B. (2018). Pharmacogenetic stimulation of neuronal activity increases myelination in an axon-specific manner. Nature Communications, 9(1), 306. https://doi.org/10.1038/s41467-017-02719-2

Moore, S., Meschkat, M., Ruhwedel, T., Tzvetanova, I. D., Trevisiol, A., Kusch, K., … Biosciences, M. (2019). A role of oligodendrocytes in information processing independent of conduction velocity.

Nave, K.-A., & Werner, H. B. (2014). Myelination of the Nervous System: Mechanisms and Functions. Annual Review of Cell and Developmental Biology, 30, 503–533. https://doi.org/10.1146/annurev-cellbio-100913-013101

Nicholson, M., Wood, R. J., Fletcher, J. L., van den Buuse, M., Murray, S. S., & Xiao, J. (2018). BDNF haploinsufficiency exerts a transient and regionally different influence upon oligodendroglial lineage cells during postnatal development. Molecular and Cellular Neuroscience, 90(November 2017), 12–21. https://doi.org/10.1016/j.mcn.2018.05.005

Nithianantharajah, J., & Hannan, A. J. (2006). Enriched environments, experience-dependent plasticity and disorders of the nervous system. Nature Reviews. Neuroscience, 7(9), 697–709. https://doi.org/10.1038/nrn1970

Okuda, H., Tatsumi, K., Makinodan, M., Yamauchi, T., Kishimoto, T., & Wanaka, A. (2009). Environmental enrichment stimulates progenitor cell proliferation in the amygdala. Journal of Neuroscience Research, 87(16), 3546–3553. https://doi.org/10.1002/jnr.22160

Perge, J. A., Niven, J. E., Mugnaini, E., Balasubramanian, V., & Sterling, P. (2012). Why do axons differ in caliber? Journal of Neuroscience, 32(2), 626–638. https://doi.org/10.1523/JNEUROSCI.4254-11.2012

Saab, A. S., Tzvetavona, I. D., Trevisiol, A., Baltan, S., Dibaj, P., Kusch, K., … Nave, K. A. (2016). Oligodendroglial NMDA Receptors Regulate Glucose Import and Axonal Energy Metabolism. Neuron, 91(1), 119–132. https://doi.org/10.1016/j.neuron.2016.05.016

Sánchez, I., Hassinger, L., Paskevich, P. A., Shine, H. D., & Nixon, R. A. (1996). Oligodendroglia regulate the regional expansion of axon caliber and local accumulation of neurofilaments during development independently of myelin formation. Journal of Neuroscience, 16(16), 5095–5105. https://doi.org/10.1523/jneurosci.16-16-05095.1996

Schain, A. J., Hill, R. A., & Grutzendler, J. (2014). Disease With Spectral Confocal Reflectance Microscopy. Nature Medicine, 20(4), 443–449. https://doi.org/10.1038/nm.3495.Label-free

Sinclair, J. L., Fischl, M. J., Alexandrova, O., Heß, M., Grothe, B., Leibold, C., & Kopp-Scheinpflug, C. (2017). Sound-evoked activity influences myelination of brainstem axons in the trapezoid body. The Journal of Neuroscience, 3728–16. https://doi.org/10.1523/JNEUROSCI.3728-16.2017

Snaidero, N., Möbius, W., Czopka, T., Hekking, L. H. P., Mathisen, C., Verkleij, D., … Simons, M. (2014). Myelin membrane wrapping of CNS axons by PI(3,4,5)P3-dependent polarized growth at the inner tongue. Cell, 156(1–2), 277–290. https://doi.org/10.1016/j.cell.2013.11.044

Snaidero, N., & Simons, M. (2017). The logistics of myelin biogenesis in the central nervous system. Glia, 65(7), 1021–1031. https://doi.org/10.1002/glia.23116

Spitzer, S. O., Sitnikov, S., Kamen, Y., Evans, K. A., Kronenberg-Versteeg, D., Dietmann, S., … Káradóttir, R. T. (2019). Oligodendrocyte Progenitor Cells Become Regionally Diverse and Heterogeneous with Age. Neuron, 0(0), 1–13. https://doi.org/10.1016/J.NEURON.2018.12.020

Stephan, R., Goellner, B., Moreno, E., Frank, C. A., Hugenschmidt, T., Genoud, C., … Pielage, J. (2015). Hierarchical microtubule organization controls axon caliber and transport and determines synaptic structure and stability. Developmental Cell, 33(1), 5–21. https://doi.org/10.1016/j.devcel.2015.02.003

Sturrock, R. R. (1980). Myelination of the Mouse Corpus Callosum. Neuropathology and Applied Neurobiology, 6(6), 415–420. https://doi.org/10.1111/j.1365-2990.1980.tb00219.x

Timmler, S., & Simons, M. (2019). Grey matter myelination. Glia, (January), 1–8. https://doi.org/10.1002/glia.23614

Venkataramani, V., Tanev, D. I., Strahle, C., Studier-Fischer, A., Fankhauser, L., Kessler, T., … Kuner, T. (2019). Glutamatergic synaptic input to glioma cells drives brain tumour progression. Nature, 573(7775), 532–538. https://doi.org/10.1038/s41586-019-1564-x

Venkatesh, H. S., Morishita, W., Geraghty, A. C., Silverbush, D., Gillespie, S. M., Arzt, M., … Monje, M. (2019). Electrical and synaptic integration of glioma into neural circuits. Nature, 573(7775), 539–545. https://doi.org/10.1038/s41586-019-1563-y

Waxman, S. G., & Sims, T. J. (1984). Specificity in central myelination: evidence for local regulation of myelin thickness. Brain Research, 292(1), 179–185. https://doi.org/10.1016/0006-8993(84)90905-3

Xiao, L., Ohayon, D., McKenzie, I. A., Sinclair-Wilson, A., Wright, J. L., Fudge, A. D., … Richardson, W. D. (2016). Rapid production of new oligodendrocytes is required in the earliest stages of motor-skill learning. Nature Neuroscience, 19(9), 1210–1217. https://doi.org/10.1038/nn.4351

Yergert, K. M., Hines, J. H., & Appel, B. (2019). Neuronal Activity Enhances mRNA Localization to Myelin Sheaths During Development, 1–30.

Yeung, M. S. Y., Zdunek, S., Bergmann, O., Bernard, S., Salehpour, M., Alkass, K., … Fris??n, J. (2014). Dynamics of oligodendrocyte generation and myelination in the human brain. Cell, 159(4), 766–774. https://doi.org/10.1016/j.cell.2014.10.011

Young, K. M., Psachoulia, K., Tripathi, R. B., Dunn, S. J., Cossell, L., Attwell, D., … Richardson, W. D. (2013). Oligodendrocyte dynamics in the healthy adult CNS: Evidence for myelin remodeling. Neuron, 77(5), 873–885. https://doi.org/10.1016/j.neuron.2013.01.006

Zhao, Y.-Y., Shi, X.-Y., Qiu, X., Lu, W., Yang, S., Li, C., … Tang, Y. (2012). Enriched Environment Increases the Myelinated Nerve Fibers of Aged Rat Corpus Callosum. The Anatomical Record: Advances in Integrative Anatomy and Evolutionary Biology, 295(6), 999–1005. https://doi.org/10.1002/ar.22446

